# piggyBac-Mediated Genomic Integration of Linear dsDNA-Based Library for Deep Mutational Scanning in Mammalian Cells

**DOI:** 10.1101/2022.01.17.476579

**Authors:** Yi Wang, Yanjie Zhao, Yifan Li, Kaili Zhang, Yan Fan, Weijun Su, Shuai Li

## Abstract

Deep mutational scanning (DMS) makes it possible to perform massively parallel quantification of the relationship between genetic variants and phenotypes of interest. However, the difficulties in introducing large variant libraries into mammalian cells greatly hinder DMS under physiological states. Here we developed two novel strategies for DMS library construction in mammalian cells, namely ‘piggyBac-*in-vitro* ligation’ and ‘piggyBac-*in-vitro* ligation-PCR’. For the first strategy, we took the ‘*in-vitro* ligation’ approach to prepare high-diversity linear dsDNAs, and integrate them into the mammalian genome with a piggyBac transposon system. For the second strategy, we further added a PCR step using the *in-vitro* ligation dsDNAs as templates, for the construction of high-content genome-integrated libraries via large-scale transfection. Both strategies could successfully establish genome-integrated EGFP-chromophore randomized libraries in HEK293T cells and enrich the green fluorescence-chromophore amino acid sequences. And we further identified a novel transcriptional activator peptide with the ‘piggyBac-*in-vitro* ligation-PCR’ strategy. Our novel strategies greatly facilitate the construction of large variant DMS library in mammalian cells, and may have great application potential in the future.

## Introduction

The rapid development and extensive application of high-throughput sequencing (HTS) technologies in the past 20 years has made it practical to yield and analyze gigabases of sequencing data at relatively low cost. It is against this background that deep mutational scanning (DMS), a massively parallel genetic assay to characterize protein sequence-function relationship, has made great progress ^[1-3]^. DMS has a wide range of applications, including protein engineering ^[4-6]^, therapeutic antibody optimization ^[7, 8]^, vaccine design ^[9-11]^, disease-associated mutation identification ^[12-16]^, and so on. In a typical DMS research design, tens or hundreds of thousands of amino acid variants are introduced into a protein (or region of a protein) of interest, and an appropriate biological assay is applied as a selection pressure, and finally HTS is performed to score each amino acid substitute according to the changes in frequency during selection, thereby achieving a high-resolution understanding of the functional consequence of each variant ^[1, 17]^.

One of the major objects of DMS is to understand the effects of protein-coding variants on humans, thus cultured mammalian cells act as a preferable platform for phenotypic screening to capture the native context, providing physiological protein folding, endogenous post-translational modifications, and also other components in associated signaling pathways at their native state ^[16, 18]^. The initial step to carry out a DMS assay is to synthesize a library of mutated variants and to introduce it into cells with proper vectors. However, the establishment of large-variant libraries in cultured mammalian cells is still challenging. Direct plasmid DNA transfection is easy to perform, but only induces transient expression of recombinant proteins. Genomic integrated libraries would be a more appropriate choice for DMS in mammalian cells. One category of approaches relies on the preparation of plasmid libraries, which are integrated into the mammalian genome via either viral-, transposon-, or recombinase-system. In these plasmid-based strategies, the initial library size can reach up to 10^9^-10^10^, which is limited by the efficiency of electro-transformation. However, the library is constantly losing its capacity during the subsequent series of steps, the plasmid amplification, extraction and transfection, resulting in a significant reduction in the ultimate library size (∼10^4^-10^5^) ^[7, 17, 19]^. And the complicated process determines these approaches to be labor intensive and time consuming. The other category of approaches is plasmid-free, mainly CRISPR/Cas9-based strategies ^[6, 7, 12, 15]^. Although there is no need to worry about the shrinkage of library size, CRISPR/Cas9-based strategies necessitate the pre-integration of landing pads into the mammalian genome, and the limited length (< 200 bp) of single-stranded oligo DNA (ssODN) using as the homology-directed repair (HDR) template also makes the insertion of large transgenes difficult. Moreover, the efficiency of CRISPR/Cas9-based strategies largely depends on the host DNA repair machinery, and it is hard to distinguish the cells undergone successful recombination from the bulk cell population.

To introduce large variant libraries into the mammalian genome in a relatively simple way for DMS study, we revised our previously developed plasmid-independent library construction strategy called ‘*in-vitro* ligation’, a method that achieves transient library expression in mammalian cells via transfection of double-stranded DNAs (dsDNAs) with intact expression cassette which is obtained by ligation of PCR-generated dsDNAs containing degenerate codons ^[20]^. We sought to improve the ‘*in-vitro* ligation’ strategy by: (i) employing a piggyBac transposon system to integrate the *in-vitro* ligation dsDNAs into the mammalian genome, and (ii) boosting the preparation yield of full-length *in-vitro* ligation dsDNAs via PCR reaction, thus facilitating large-scale transfection. Here we describe these improved strategies, which we refer to as ‘piggyBac-*in-vitro* ligation’ and ‘piggyBac-*in-vitro* ligation-PCR’ strategies, and validate their applications via the identification of green fluorescence-chromophore amino acid sequences and also a novel transcriptional activator peptide.

## Materials and Methods

### Cell culture

The HEK293T cell line was obtained from the Type Culture Collection of the Chinese Academy of Sciences (Shanghai, China) and was maintained in Dulbecco’s Modified Eagle’s medium (DMEM) (Corning, NY, USA) containing 10% fetal bovine serum (FBS) (Corning) and 1% penicillin-streptomycin solution (10,000 U/ml, Thermo Fisher Scientific, Waltham, MA, USA) at 37 °C with 5% CO_2_.

### Plasmid construction

The plasmids used as PCR templates (PuroR-template-plasmid, EGFP-chromophore^del^-library-template-plasmid and GAL4-XTEN-library-template-plasmid) and EGFP-IRES-mCherry plasmid were constructed by VectorBuilder (VectorBuilder, Guangzhou, China) (for the sequences of plasmids, please refer to the **Supporting Information**). Plasmids coding EGFP-mutants were generated from EGFP-IRES-mCherry plasmid using TaKaRa MutanBEST Kit (TaKaRa, Kusatsu, Japan). pG5egfp reporter vector was modified from the pG5luc vector (GeneBank Accession Number AF264724) as we mentioned before ^[21]^. pBIND-XTEN-VP16, pBIND-XTEN-VP32, pBIND-XTEN-VP64, pBIND-XTEN-TAP1, pBIND-XTEN-(TAP1)_2_ and pBIND-XTEN-TAP1-VP16 plasmids were constructed using the pBIND vector (GeneBank Accession Number AF264722) as their backbones. Specifically, sequences encoding XTEN-peptide were synthesized and cloned between XbaI and KpnI sites.

### Synthesis of dsDNAs for library construction

High-fidelity KOD-Plus-Neo DNA polymerase (Toyobo, Osaka, Japan) was applied to perform PCR amplification. All PCR templates and the corresponding primers used in this study are listed in **Supplementary Table 1**.

For the construction of piggyBac-*in-vitro* ligation libraries, the PCR products were purified after agarose electrophoresis (using Universal DNA purification kit, TIANGEN, Beijing, China), and their concentrations were determined by the OD 260 value by NanoDrop (Thermo Fisher Scientific). The upstream and downstream PCR products were digested by BssSI restriction enzyme (New England Biolabs, Ipswich, MA, USA). After another round of agarose electrophoresis and gel purification, the upstream and downstream digested dsDNA were ligated to each other by T4 DNA ligase (New England Biolabs). To unbind the ligase from DNA, the ligation samples were pre-incubated with 6× DNA Loading Dye & SDS Solution (Thermo Fisher Scientific) at 65°C for 10 minutes and chilled on ice before the agarose electrophoresis. Then the *in-vitro* ligation products were further purified for the following transfection. To avoid UV-induced DNA damage, Blue Light Gel Imager (Sangon Biotech, Shanghai, China) was used for DNA visualization.

For the construction of piggyBac-*in-vitro* ligation-PCR libraries, 1 or 10 ng full-length dsDNAs generated by the *in-vitro* ligation strategy were used as templates for each 50 μl standard PCR reaction system to generate *in-vitro* ligation-PCR (1 ng) or *in-vitro* ligation-PCR (10 ng) library dsDNAs respectively. Numbers of PCR cycles for each library were determined to ensure that the amplification was in the exponential phase.

### Cell transfection, resistance selection, and colony formation assay

For the test of piggyBac-mediated genomic integration of *in-vitro* ligation dsDNAs, HEK293T cells were plated in 6-well plates the day before transfection, and reached a 70∼80 % confluency on the day of transfection. *in-vitro* ligation dsDNAs (1 μg/well), PCR products (1 μg/well), or plasmids (1 μg/well) were respectively co-transfected with hyperactive piggyBac transposase expression vector (1 μg/well) into HEK293T cells using Lipofectamine 2000 (Thermo Fisher Scientific). Forty-eight hours after transfection, cells were 1,000 times diluted, and a 10-day selection with puromycin (Selleck Chemicals, Houston, TX, USA) was performed to select the cells with successful genomic integration. Ten days later, the crystal violet staining was performed with the kit from Beyotime Biotechnology (Shanghai, China) according to the manufacturer’s instruction, and the representative pictures were taken.

For the transfection of piggyBac-*in-vitro* ligation library, HEK293T cells were plated in 6-well plates the day before transfection, and reached a 70∼80% confluency on the day of transfection. piggyBac-*in-vitro* ligation library dsDNAs (1 μg/well) were co-transfected with hyperactive piggyBac transposase expression vector (1 μg/well) into HEK293T cells using Lipofectamine 2000. For the transfection of piggyBac-*in-vitro* ligation-PCR library, HEK293T cells were plated in 10-cm dishes the day before transfection, and reached a 70∼80% confluency on the day of transfection. Library dsDNAs (6 μg/dish) and hyperactive piggyBac transposase expression vector (6 μg/dish) were co-transfected into HEK293T cells using Lipofectamine 2000. Forty-eight hours after transfection, a 10-day selection with puromycin was performed.

For the identification of novel transcriptional activator peptide, HEK293T cells were plated in 10-cm dishes the day before transfection, and reached a 70-80% confluency on the day of transfection. Library dsDNAs with the promoter-GAL4-XTEN-<NNK>_12_-PAS expression cassette (6 μg/dish) and hyperactive piggyBac transposase expression vector (6 μg/dish) were co-transfected into HEK293T cells using Lipofectamine 2000. Forty-eight hours after transfection, a 5-day selection with puromycin was performed. The obtained cells with successful genomic integration were re-seeded into eight 10-cm dishes and transfected with pG5egfp vector (6 μg/dish). Forty-eight hours after transfection, cells were harvested for the following FACS sorting.

For the verification of the transcriptional activation potential of TAP1, HEK293T cells were plated in 24-well plates the day before transfection, and reached a 70∼80% confluency on the day of transfection. pBIND-XTEN-vector, pBIND-XTEN-VP16, pBIND-XTEN-VP32, pBIND-XTEN-VP64, pBIND-XTEN-TAP1, pBIND-XTEN-(TAP1)_2_ or pBIND-XTEN-TAP1-VP16 plasmids (250ng/well) were respectively co-transfected with pG5luc plasmid (250ng/well) into HEK293T cells using Lipofectamine 2000. Forty-eight hours after transfection, the cells were harvested for the following dual-luciferase assay.

### Fluorescence activated cell sorting (FACS) and analytical flow cytometry

HEK293T cells with library genomic integration were digested with trypsin and re-suspended with PBS. Cells with green fluorescence or blue fluorescence were harvested using a BD FACSAria™ II Cell Sorter (BD Biosciences, Bedford, MA, USA) for the following experiments.

Analytical flow cytometry was performed with a CytoFLEX LX flow cytometer (Beckman Coulter Life Sciences, Indianapolis, IN, USA), and the results were analyzed with FlowJo software (FLOWJO, Ashland, OR, USA).

### High-throughput sequencing (HTS) and data processing

Genomic DNAs were extracted with a TaKaRa MiniBEST Universal Genomic DNA Extraction Kit Ver. 5.0 (TaKaRa). High-fidelity KOD-Plus-Neo DNA polymerase was used to amplify the sequences containing the degenerate codons. Library construction and high-throughput sequencing were conducted by Novogene (Beijing, China). The PCR products were subjected to sequencing library construction using the TruSeq^®^ DNA PCR-Free Sample Preparation Kit (Illumina, San Diego, CA, USA). The libraries were loaded on a NovaSeq 6000 system (Illumina) to generate PE-150 reads. Clean reads were merged using the FLASH (Fast Length Adjustment of SHort reads) software (v1.2.11). And in-house Perl scripts were developed to analyze the amino acid sequences of libraries.

The read counts of all possible amino-acid combinations in each library were first normalized as RPM (read per million reads) values. Then the RPM values from the initial and selected libraries are used to calculate the fold of enrichment for each amino-acid combination.

### Dual luciferase reporter assay

Dual reporter activities were measured using the Dual Luciferase Reporter Assay System (Promega, Madison, WI, USA) according to the manufacturer’s instructions.

### Statistics

All data are presented as mean ± SD unless otherwise stated. Differences were assessed by two-tailed Student’s *t*-test using GraphPad software (GraphPad Software, San Diego, CA, USA). *p* < 0.05 was considered to be statistically significant.

## Results

### Genomic integration of *in-vitro* ligation dsDNAs via a piggyBac transposon system

In our previous study, we developed a plasmid-independent ‘*in-vitro* ligation’ strategy to introduce DNA-encoded high-diversity nanobody libraries into mammalian cells ^[20]^. Briefly, PCR-generated upstream and downstream dsDNAs containing random (NNK) CDR sequences were digested by BssSI restriction enzyme and ligated by T4 DNA ligase to generate dsDNAs with the intact nanobody expression cassette. The obtained full-length dsDNAs were transfected into HEK293T cells for nanobody library transient expression. This strategy can generate over a million different nanobody sequences as revealed by HTS. In the present study, to retrofit this strategy for DMS library construction in mammalian cells, we further employ a piggyBac transposon system to integrate the *in-vitro* ligation dsDNAs into the mammalian genome.

The piggyBac transposon system can efficiently insert foreign DNA into the mammalian genome via a non-viral route ^[22]^. First, we designed a colony formation assay to assess the feasibility of piggyBac-mediated genomic integration of *in-vitro* ligation dsDNAs. We added the piggyBac transposon-specific inverted terminal repeat sequences (ITRs) to both the 5’ terminus of the upstream dsDNAs and the 3’ terminus of the downstream dsDNAs. And we employed T4 DNA ligase to ligate the BssSI-digested upstream and downstream dsDNAs to obtain an intact puromycin resistance gene expression cassette **(Figure 1A)**. The *in-vitro* ligation dsDNAs and a hyperactive piggyBac transposase expression vector ^[23]^ were co-transfected into HEK293T cells. Forty-eight hours later, the transfected cells were 1,000 times diluted and cultured with puromycin. After a 10-day selection, the surviving puro-resistant cell clones were stained with crystal violet **(Figure 1B)**. As shown in **Figure 1C**, we finally harvested 2.9×10^4^ puro-resistant clones from one single well (6-well plate) of HEK293T cells transfected with 1 µg *in-vitro* ligation dsDNAs. As a comparison, transfection with PCR products (namely PuroR PCR products) or plasmids (PuroR-template-plasmid) containing puro-resistance gene expression cassette and ITR sequences led to the formation of 1.98×10^5^ or 2.03×10^5^ puro-resistant cell clones per well respectively **(Figure 1C)**. So far, we have proved that the piggyBac transposon system could successfully mediate the integration of *in-vitro* ligation dsDNAs into the mammalian genome, and we named this strategy as ‘piggyBac-*in-vitro* ligation’.

**Figure 1.**
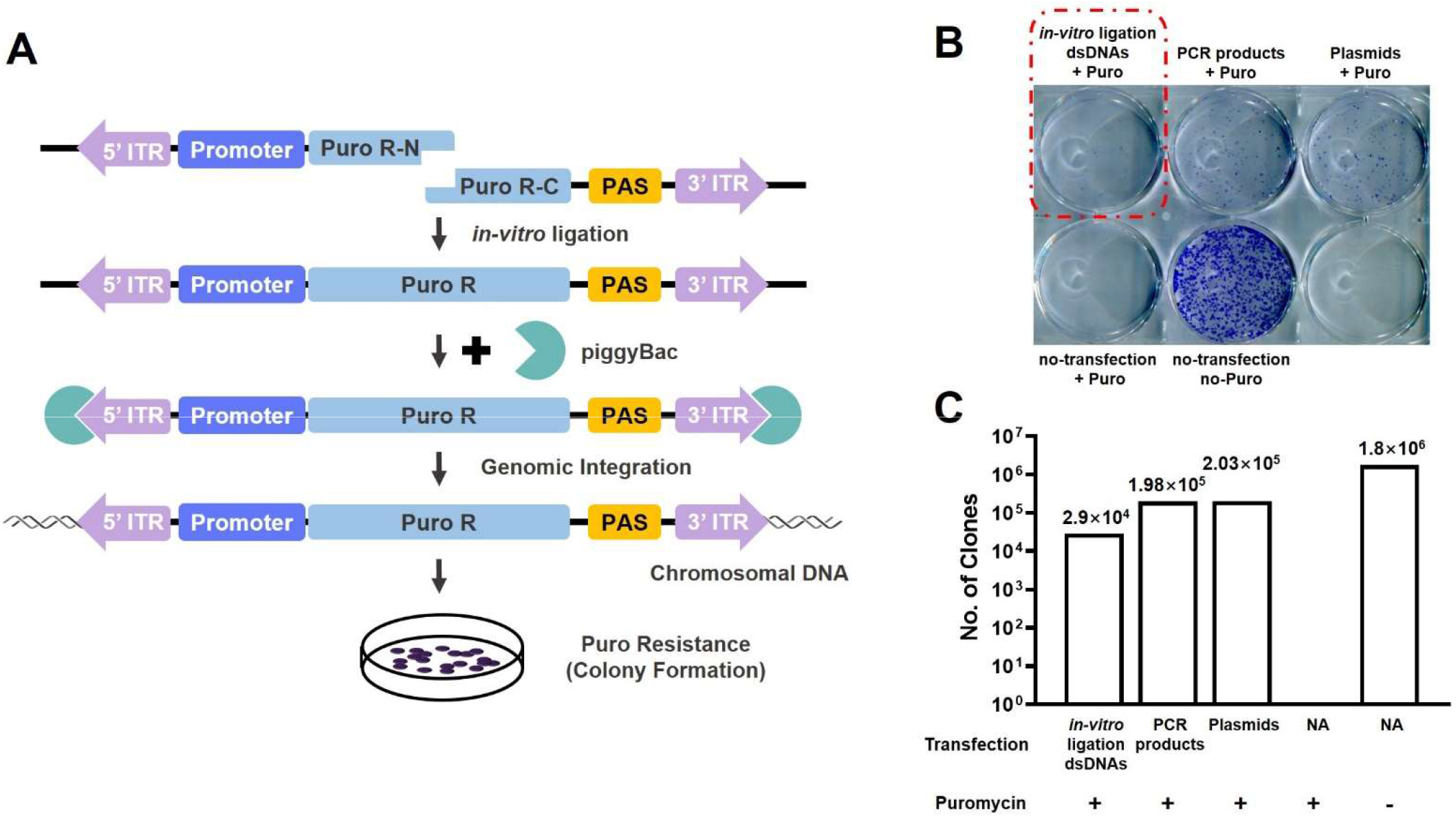
piggyBac transposon system-mediated genomic integration of *in-vitro* ligation dsDNAs. **(A)** Schematic representation of the piggyBac transposon system-mediated *in-vitro* ligation dsDNAs genomic integration strategy. Both the upstream dsDNAs with 5’ ITR and the downstream dsDNAs with 3’ ITR were digested with BssSI restriction enzyme, and were ligated with each other by T4 DNA ligase. The obtained full-length dsDNAs were with an intact puromycin resistance gene expression cassette and ITRs at both termini, and were co-transfected with a piggyBac transposase expression vector into mammalian cells. The puro-resistance gene expression cassette was integrated into the genome by the piggyBac transposon system, and endowed puromycin resistance property to the cells. **(B-C)** Colony formation assay to assess the genomic integration efficiency of *in-vitro* ligation dsDNAs by the piggyBac transposon system. The *in-vitro* ligation dsDNAs containing puro-resistance gene expression cassette and ITR sequences were co-transfected into HEK293T cells with a piggyBac transposase expression vector. Forty-eight hours later, the transfected cells were 1,000 times diluted and were cultured with puromycin. Ten days later, the puro-resistance cell clones were stained with crystal violet. The PCR product and the plasmid containing the intact puro-resistance gene expression cassette and the bilateral ITRs were used as controls. A representative image of crystal violet staining is shown in **(B)** and the statistics of the clone number in each well (the clone number in the corresponding well in **(B)** multiply by 1,000) are presented in **(C)**.

### Identification of chromophore amino acid sequences by DMS with ‘piggyBac*-in-vitro* ligation’ strategy in mammalian cells

Next, we tested the feasibility of*’*piggyBac-*in-vitro* ligation’ strategy in DMS in mammalian cells by constructing genome-integrated EGFP-chromophore-randomized libraries. Green fluoresecent protein (GFP) is widely used in a myriad of modern biological researches, providing a screenable phenotype by FACS ^[24, 25]^. The chromophore is encoded by the primary amino acid sequence (S65-Y66-G67 for *A. victoria* GFP (wtGFP) and T65-Y66-G67 for enhanced GFP (EGFP)), and there is no requirement for cofactors or external enzyme components for chromophore formation ^[26]^. The leucine at position 64 in EGFP is used to replace the phenylalanine in wtGFP, which improves the efficiency of protein maturation at 37°C ^[26]^. Here, we generated *in-vitro* ligation EGFP-chromophore-randomized libraries by NNK degeneracy, integrated them into the HEK293T cell genome with a piggyBac transposon system, and expected to enrich the sequences coding green fluorescence-generating proteins by FACS.

As shown in **Figure 2A**, the 2NNK (corresponding to EGFP T65-Y66) or 3NNK (corresponding to EGFP T65-Y66-G67) codons were incorporated into the downstream dsDNAs via PCR reaction with degenerate primers. And the full-length EGFP-chromophore-randomized (2NNK or 3NNK) dsDNAs with ITRs at both termini were generated via *in-vitro* ligation strategy, and were co-transfected with a piggyBac transposase expression vector into HEK293T cells. For the construction of each library, the transfection was performed in one single well (6-well plate). Forty-eight hours after transfection, a 10-day puromycin selection was performed, followed by FACS sorting of the cells with green fluorescence. Green fluorescent cells were collected and cultured for the subsequent round of sorting.

**Figure 2.**
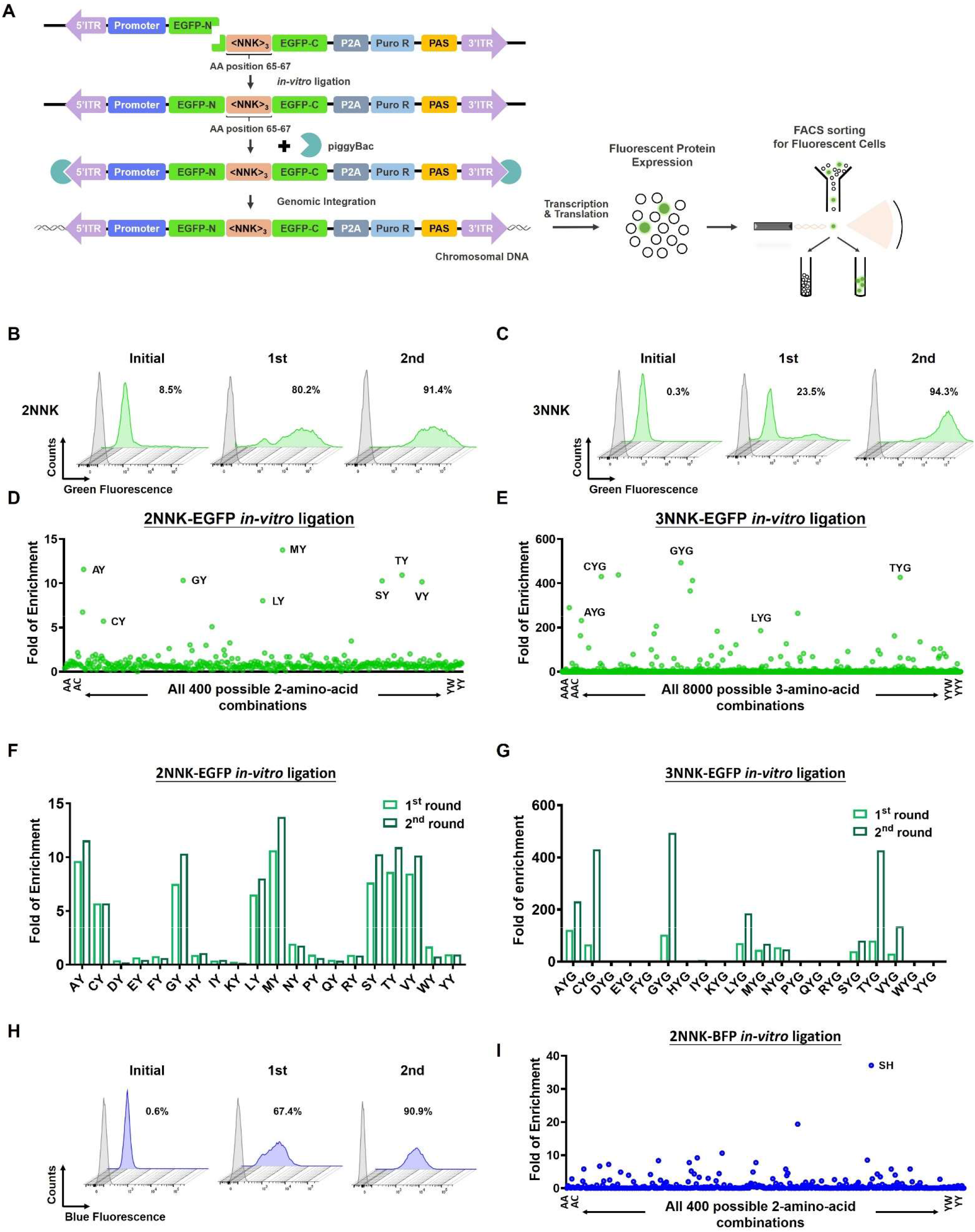
The application of ‘piggyBac-*in-vitro* ligation’ strategy in the identification of chromophore amino acid sequences. **(A)** Schematic representation of ‘piggyBac-*in-vitro* ligation’ strategy for the identification of chromophore amino acid sequences. In the downstream dsDNAs, EGFP chromophore-randomized sequence (corresponding to T65-Y66-G67) was introduced by NNK degenerate codons, and the puro-resistance gene was linked to the downstream of EGFP coding region via a P2A element. The full-length dsDNAs with chromophore-randomized EGFP expression cassette and bilateral ITR sequences were generated via the *in-vitro* ligation strategy. After co-transfection with a piggyBac transposase expression vector into mammalian cells, the chromophore-randomized EGFP expression cassette was integrated into the genome. Forty-eight hours after transfection, the cells were cultured with puromycin for 10 days. Rounds of FACS sorting were performed to harvest the fluorescent cells, and HTS was performed to analyze the chromophore amino acid sequences. **(B-C)** The ratios of green fluorescent cells after each round of FACS sorting in 2NNK-EGFP *in-vitro* ligation library **(B)** and 3NNK-EGFP *in-vitro* ligation library **(C). (D-E)** The fold of enrichment of each possible 2-amino-acid combination in 2NNK-EGFP *in-vitro* ligation library **(D)** and each possible 3-amino-acid combination in 3NNK-EGFP *in-vitro* ligation library **(E)** after 2 rounds of FACS sorting of green fluorescent cells. After 2 rounds of sorting, the DNA sequences containing the degenerate codons were PCR-amplified and subjected to HTS. The read count of each possible amino acid combination was normalized as RPM value, and the fold of enrichment was calculated. The amino-acid combinations are alphabetically ordered on X-axis. **(F-G)** The fold of enrichment of each X65-Y66-pattern 2-amino-acid combination in 2NNK-EGFP *in-vitro* ligation library **(F)** and each X65-Y66-G67-pattern 3-amino-acid combination in 3NNK-EGFP *in-vitro* ligation library **(G)** after each round of FACS sorting. **(H)** The ratios of blue fluorescent cells after each round of FACS sorting in 2NNK-BFP *in-vitro* ligation library. **(I)** The fold of enrichment of each possible 2-amino-acid combination in 2NNK-BFP *in-vitro* ligation library after 2 rounds of FACS sorting of blue fluorescent cells. The amino acid combinations are alphabetically ordered on X-axis.

After 2 rounds of FACS sorting, the ratio of green fluorescent cells reached 91.4% for 2NNK-EGFP *in-vitro* ligation library **(Figure 2B)** and 94.3% for 3NNK-EGFP *in-vitro* ligation library **(Figure 2C)**. The genomic DNAs of the initial puro-selected cells and the cells after each round of FACS sorting were extracted, and the DNA sequences containing the degenerate codons were analyzed by HTS. We counted the RPM (read per million reads) for all 400 possible 2-amino-acid combinations in 2NNK-EGFP *in-vitro* ligation library and 8000 possible 3-amino-acid combinations in 3NNK-EGFP *in-vitro* ligation library, and calculated the fold of enrichment after each round of FACS sorting. For the 2NNK-EGFP *in-vitro* ligation library, 8 out of the top 10 enriched 2-amino-acid combinations showed a conserved X65-Y66 pattern **(Figure 2D&F, Supplementary Table 2)**. For the 3NNK-EGFP *in-vitro* ligation library, 5 out of the top 15 enriched 3-amino-acid combinations showed a conserved X65-Y66-G67 pattern **(Figure 2E&G, Supplementary Table 3)**.

As one of the earliest color variants derived from GFP, blue fluorescent protein (BFP) contains a Y66H substitution ^[26]^. Accordingly, we also tried to collect the cells with blue fluorescence from the HEK293T cells transfected with the 2NNK-EGFP *in-vitro* ligation library following the same strategy. The ratio of blue fluorescent cells was enhanced from 0.6% to 90.9% after 2 rounds of FACS sorting **(Figure 2H)**. And the HTS results revealed that the fold of enrichment of the original BFP-chromophore amino acid sequence S65-H66 ranked 1^st^ among all 400 possible 2-amino-acid combinations after 2 rounds of selection **(Figure 2I, Supplementary Table 4)** ^[27]^.

### Development of ‘piggyBac-*in-vitro* ligation-PCR’ strategy for genome-integrated high-content DMS library construction

So far, we have proved the applicability of ‘piggyBac-*in-vitro* ligation’ strategy to DMS library construction. But there is an inherent flaw of this strategy: since the synthesis requires multi-step enzymatic reactions, it is difficult to prepare *in-vitro* ligation dsDNAs in large quantities, which hinders the establishment of genome-integrated high-content DMS libraries via large-scale transfection. Thus, we further supplemented a PCR step to this strategy to amplify the full-length *in-vitro* ligation dsDNAs, and named it as ‘piggyBac-*in-vitro* ligation-PCR’ strategy. We also tested its feasibility by tentative identification of green fluorescence-chromophore amino acid sequences **(Figure 3A)**.

**Figure 3.**
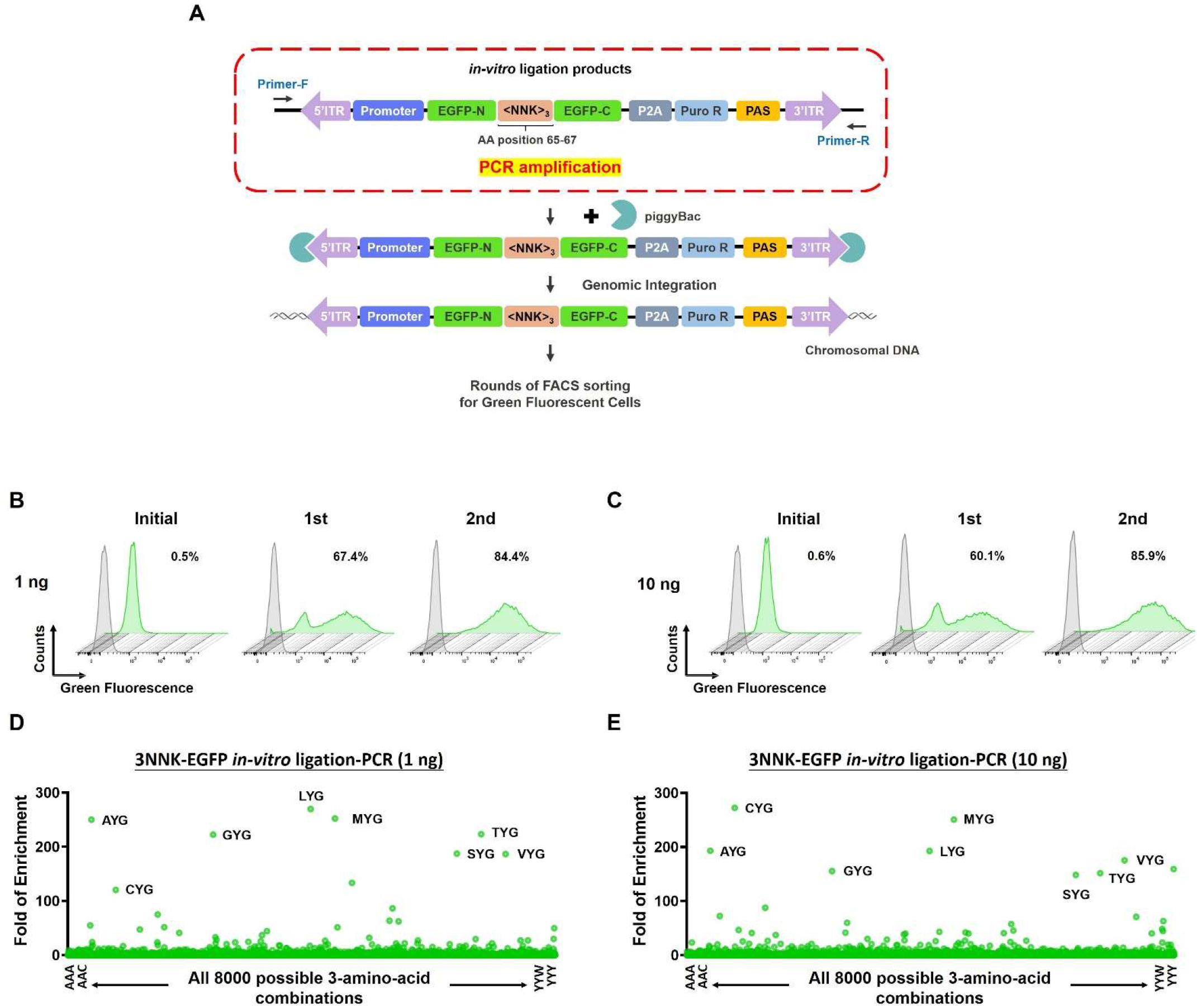
The application of ‘piggyBac-*in-vitro* ligation-PCR’ strategy for the identification of green fluorescence-chromophore amino acid sequences. **(A)** Schematic representation of ‘piggyBac-*in-vitro* ligation-PCR’ strategy for the establishment of genome-integrated 3NNK-EGFP *in-vitro* ligation-PCR library. On the basis of the ‘piggyBac-*in-vitro* ligation’ strategy, a PCR step (indicated by the red dashed line) was added to amplify the full-length *in-vitro* ligation dsDNAs. **(B-C)** The ratios of green fluorescent cells after each round of FACS sorting in the genome-integrated 3NNK-EGFP *in-vitro* ligation-PCR libraries. Different amounts of *in-vitro* ligation dsDNA templates were used for PCR reaction in the construction of DMS libraries as indicated. **(D-E)** The fold of enrichment of each possible 3-amino-acid combination after 2 rounds of FACS sorting of green fluorescent cells in the genome-integrated 3NNK-EGFP *in-vitro* ligation-PCR libraries. Different amounts of *in-vitro* ligation dsDNA templates were used for PCR reaction in the construction of DMS libraries as indicated. The amino acid combinations are alphabetically ordered on X-axis.

Specifically, the full-length EGFP-chromophore-randomized (3NNK) dsDNAs with ITRs at both termini generated by the *in-vitro* ligation strategy were used as templates in PCR for amplification. Different amounts of *in-vitro* ligation dsDNA templates were used in the PCR system (1 or 10 ng dsDNAs/50 µl PCR system). The PCR products were then integrated into the HEK293T cell genome via the piggyBac transposon system, thereby generating genome-integrated EGFP-chromophore-randomized (3NNK) libraries **(Figure 3A)**. For the construction of each library, one 10-cm dish of HEK293T cells were transfected. Two successive rounds of FACS sorting enhanced the ratio of green fluorescent cells to around 85% in both high (10 ng) and low (1 ng) amount of PCR template group **(Figure 3BC)**. And the HTS analysis revealed the same eight 3-amino-acid combinations (AYG, CYG, GYG, LYG, MYG, SYG, TYG, and VYG) conforming to the X65-Y66-G67 pattern among the top 10 FACS-enriched sequences in both libraries **(Figure 3DE, Supplementary Figure 1, Supplementary Table 5-6)**.

This ‘piggyBac-*in-vitro* ligation-PCR’ strategy helps to establish genome-integrated high-content DMS libraries via large-scale transfection, thus theoretically enables the screening for extremely rare phenotypes. To demonstrate it, we diluted the PCR-amplified (10 ng template per 50 µl PCR system) EGFP-chromophore-randomized (3NNK) dsDNAs 100 times with EGFP-chromophore-deleted dsDNAs to artificially simulate a relatively rare phenotype, as shown in **Figure 4A**. Except for the deletion of chromophore-coding sequence (9 nt), the EGFP-chromophore-deleted dsDNAs have the same sequence with EGFP-chromophore-randomized (3NNK) dsDNAs. The mixture of dsDNAs were transfected into two 10-cm dish-seeded HEK293T cells, and a piggyBac transposon system mediated their genomic integration. Two rounds of FACS sorting were performed to enrich green fluorescent cells **(Figure 4B)**. After rounds of enrichment, the HTS analysis revealed eight 3-amino-acid combinations conforming to the X65-Y66-G67 pattern (SYG, TYG, VYG, GYG, LYG, AYG, MYG, CYG) among the top 15 hits **(Figure 4C)**, which were exactly the same as the eight X65-Y66-G67-pattern sequences screened out from the undiluted 3NNK-EGFP *in-vitro* ligation-PCR library **(Figure 3E)**.

**Figure 4.**
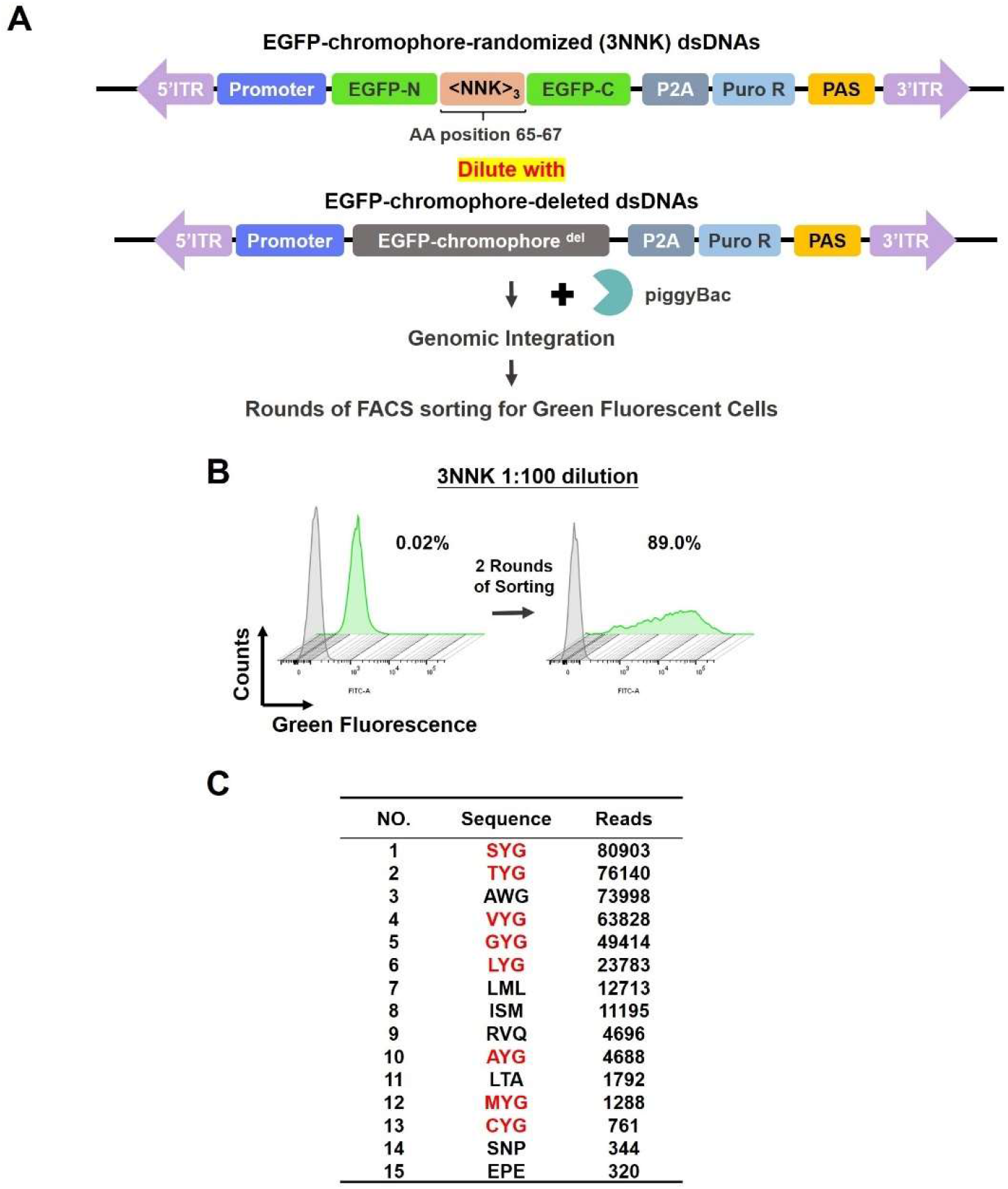
Test of the DMS library constructed by ‘piggyBac-*in-vitro* ligation-PCR’ strategy in the screening for relatively rare phenotype. **(A)** Schematic diagram of the mixture of PCR-amplified EGFP-chromophore-randomized (3NNK) dsDNAs with EGFP-chromophore-deleted dsDNAs for the establishment of the 1:100 diluted 3NNK-EGFP *in-vitro* ligation-PCR library. **(B)** The ratio of green fluorescent cells after 2 rounds of FACS sorting in the 1:100 diluted 3NNK-EGFP *in-vitro* ligation-PCR library. **(C)** The top fifteen 3-amino-acid combinations enriched by 2 rounds of FACS sorting for green fluorescent cells from the 1:100 diluted 3NNK-EGFP *in-vitro* ligation-PCR library, as revealed by HTS analysis. The combinations conforming to X65-Y66-G67 pattern are labeled in red.

### Assessment of green fluorescence in EGFP-mutants

Our above results showed high repetition rate among the top enriched sequences with the 3NNK-EGFP *in-vitro* ligation library **(Figure 2E)**, the 3NNK-EGFP *in-vitro* ligation-PCR library **(Figure 3DE)**, and the diluted 3NNK-EGFP *in-vitro* ligation-PCR library by HTS analysis **(Figure 4C)**. Most of the top hits presented a conserved X65-Y66-G67 pattern, which hints the indispensability of the tyrosine at AA position 66 and the glycine at AA position 67 for green fluorescence generation. These results are highly consistent with the reported critical roles of Y66 in GFP absorption-emission spectral profile maintenance and G67 in imidazolinone ring formation for chromophore maturation ^[26, 28]^. For further evidence, we constructed two EGFP-mutant plasmids in which the tyrosine at position 66 or the glycine at position 67 was mutated to alanine respectively (named as TAG or TYA), while an IRES-mCherry sequence was fused to the downstream of EGFP framework, serving as a reference ^[17]^ **(Figure 5A)**. As we expected, alanine mutation at either position 66 or 67 extinguished the green fluorescence by EGFP (T65-Y66-G67) **(Figure 5A)**.

**Figure 5.**
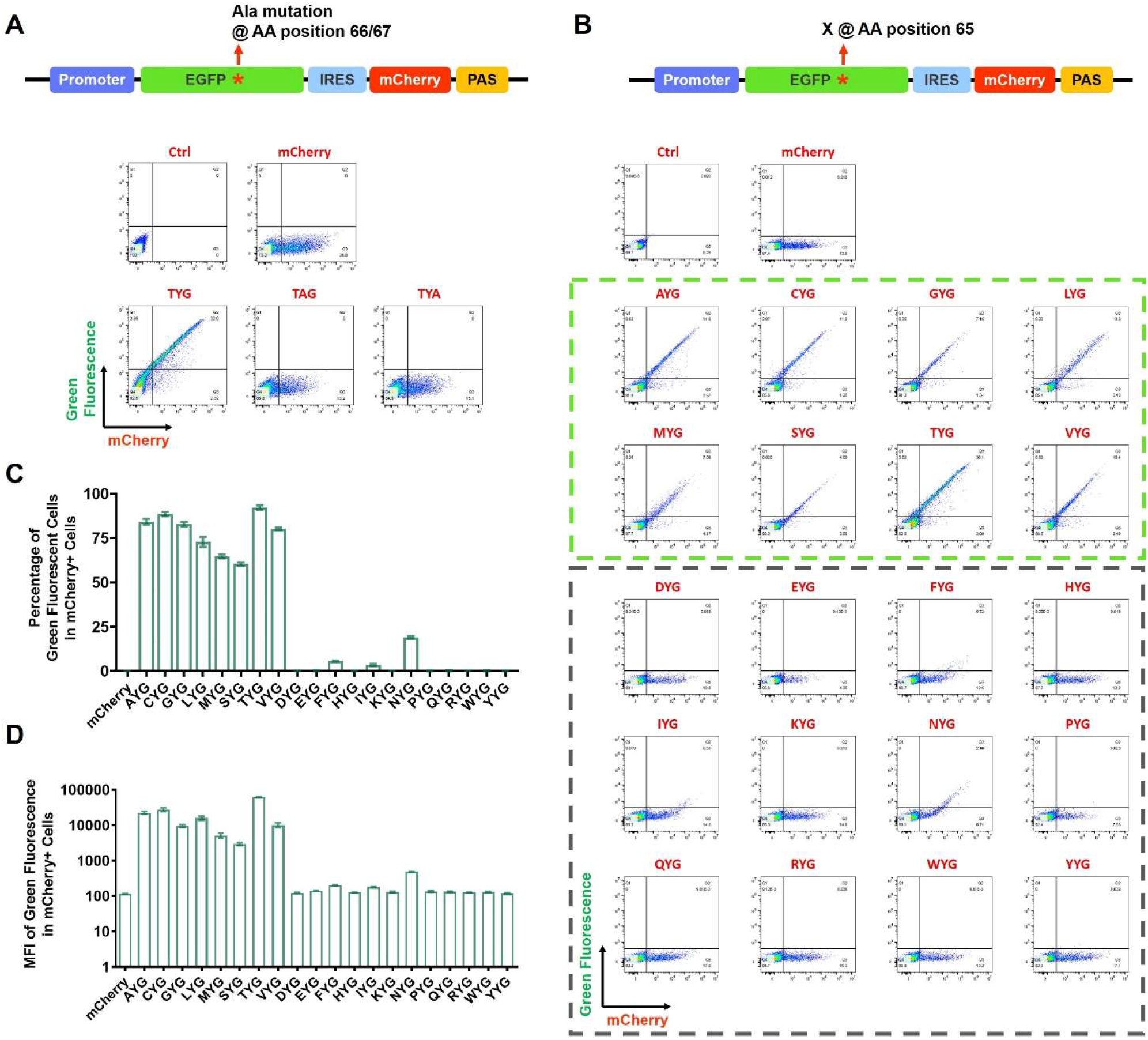
Assessment of green fluorescence in EGFP-mutants on AA position 65, 66 and 67. **(A)** Detection of green fluorescence by Y66A and G67A EGFP-mutants via FACS. Two plasmids encoding Y66A and G67A EGFP-mutants were constructed, in which mCherry was used as a reference. EGFP-mutant plasmids were transfected into HEK293T cells. Forty-eight hours later, the cells were harvested and detected for green fluorescence and mCherry expression by FACS. Representative dot plots for each mutant are present. **(B-D)** Detection of green fluorescence by EGFP-mutants on AA position 65 via FACS. Twenty plasmids encoding EGFP-mutants with all possible canonical amino acids at position 65 were constructed, in which mCherry was used as a reference. EGFP-mutant plasmids were transfected into HEK293T cells. Forty-eight hours later, the cells were harvested and detected for green fluorescence and mCherry expression by FACS. Representative dot plots for each mutant are present in **(B)**. Percentages of green fluorescent cells in mCherry^+^ cells for each mutant are shown in **(C)**. Mean fluorescence intensity (MFI) of green fluorescence in mCherry^+^ cells for each mutant are shown in **(D)**. All data are displayed as the mean ± SD (n = 3).

To further systematically analyze the impact of various amino acids at AA position 65 on green fluorescence generation, we constructed EGFP-mutant plasmids encoding all 20 possible canonical amino acids at position 65, and employed mCherry as a reference **(Figure 5B)**. The results of analytical flow cytometry showed that the eight EGFP-mutants which encode the Ala, Cys, Gly, Leu, Met, Ser, Thr or Val at AA position 65, but not the other 12 mutants, could generate detectable green fluorescence under 488-nm laser excitation **(Figure 5B-D)**, which was highly consistent with the above HTS results. These results fully proved the applicability of our novel strategies in the DMS library construction in mammalian cells.

### Identification of a novel transcriptional activator peptide by ‘piggyBac-*in-vitro* ligation-PCR’ strategy

To further demonstrate the applicability of our novel strategy in functional screening, we sought to create a workflow for novel transcriptional activator peptide identification in mammalian cells. We established a 12-amino-acid peptide library with ‘piggyBac-*in-vitro* ligation-PCR’ strategy, linked it to the downstream of GAL4 DNA-binding domain via a 16-amino-acid XTEN linker. The promoter-GAL4-XTEN-<NNK>_12_-PAS expression cassette was integrated into the genome of HEK293T cells via the piggyBac transposon system, and the generated 12-amino-acid peptides were tested in transient transfection with a pG5egfp reporter vector containing five GAL4-binding sites in a promoter position of EGFP **(Figure 6A)**. After 2 rounds of FACS sorting, EGFP positive cells were harvested for genomic DNA extraction and the following HTS analysis. Top 10 possible transcriptional activator peptides with the most read counts were listed in **Figure 6B**. Among them, the read count number of peptide LIFTYMDNFDWW ranking 1^st^ (hereafter called TAP1) was much higher than that of the rests **(Figure 6B)**. Furthermore, we found that there is a characteristic sequence motif (DNFDW) containing hydrophobic residue (Phe, F) flanking by carbonyl-containing amino acids, the analog of which can be found in a series of well-known transcriptional activator domains, including the VP16 domain (residues 437-447, DAL<u>DDFDL</u>DML) ^[29]^. Thus, we cloned the coding sequence of TAP1 into pBIND vector coding GAL4 DNA-binding domain and assessed its transcriptional activation potential in transient transfection with pG5luc containing five GAL4 binding sites upstream of a minimal TATA box and luciferase CDS, and the VP16 domain was used as a positive control. The transcriptional activation ability of TAP1 is greater than that of the VP16 domain in this system **(Figure 6C)**. We further investigated the transcriptional activation potential of bipartite peptides (TAP1)_2_ and TAP1-VP16. TAP1-VP16 has a slightly lower transcriptional activity to VP32, while unexpectedly, two identical TAP1 copies as a tandem repeat has little transcriptional activation ability **(Supplementary Figure 2)**.

**Figure 6.**
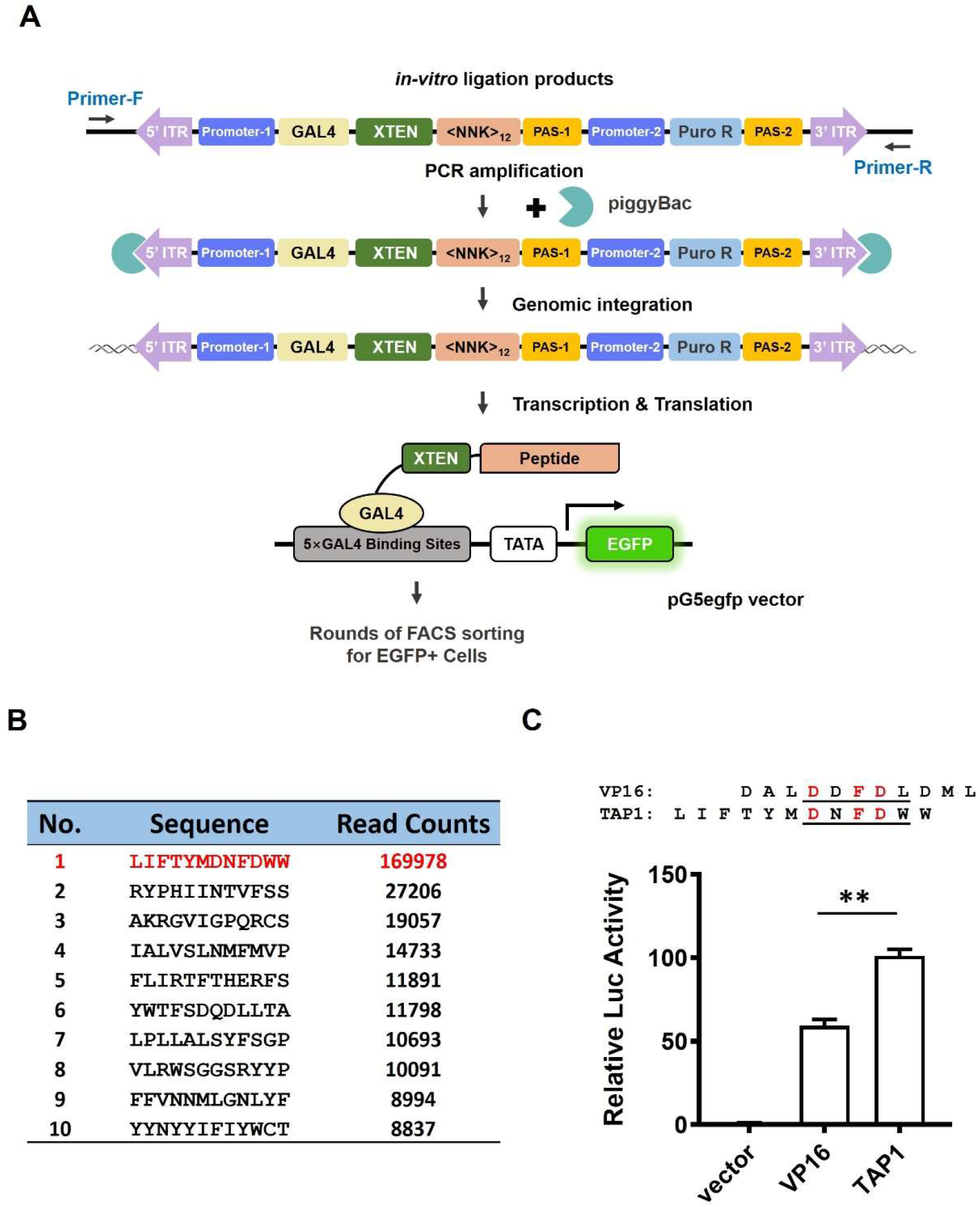
Screening of transcriptional activator peptides using ‘piggyBac-*in-vitro* ligation-PCR’ strategy. **(A)** Schematic representation of transcriptional activator peptide screening workflow using ‘piggyBac-*in-vitro* ligation-PCR’ strategy. *In-vitro* ligation products containing the promoter-GAL4-XTEN-<NNK>_12_-PAS expression cassette flanked by ITRs were PCR-amplified, and the PCR products were co-transfected with a piggyBac transposase expression vector into mammalian cells. Forty-eight hours later, a 7-day puromycin selection was performed, thus HEK293T cells with genomic integration of promoter-GAL4-XTEN-<NNK>12-PAS expression cassette were harvested. A pG5egfp reporter vector containing five GAL4-binding sites in a promoter position of EGFP was further transfected into the cells. The interaction between GAL4-binding sites in the pG5egfp vector and the GAL4 DNA-binding domain pulled the 12-amino-acid peptides to the proximity of TATA box, and the 12-amino-acid peptides with transcriptional activity initiated EGFP reporter expression. Rounds of FACS sorting were performed to enrich the EGFP positive cells for further HTS analysis. **(B)** The sequences and read count numbers of the top ten 12-amino-acid transcriptional activator peptide candidates revealed by HTS analysis. **(C)** Transcriptional activation potential of peptide TAP1. The coding sequence of TAP1 was cloned into pBIND vector coding GAL4 DNA-binding domain (namely pBIND-XTEN-TAP1), and co-transfected with pG5luc vector containing five GAL4 binding sites upstream of a minimal TATA box and luciferase CDS into HEK293T cells. The transcriptional activity of TAP1 was assessed by the relative Luc activity (Fluc/Rluc) measured by the dual luciferase reporter assay, and compared to that of VP16 domain. All data are displayed as the mean ± SD (n = 4). \*\**p* < 0.01.

## Discussion

In this study, we have described two novel strategies for DMS assay in mammalian cells, namely ‘piggyBac-*in-vitro* ligation’ and ‘piggyBac-*in-vitro* ligation-PCR’, and demonstrated their capability. These strategies were developed on the basis of ‘*in-vitro* ligation’ strategy we reported previously ^[20]^, with the help of piggyBac transposon system for genomic integration and PCR reaction for mass preparation of dsDNAs for transfection. With either of these strategies, we successfully established genome-integrated EGFP-chromophore randomized library in HEK293T cells and enriched the green fluorescence-chromophore amino acid sequences. Moreover, we also identified a brand new 12-amino-acid transcriptional activator peptide with the ‘piggyBac-*in-vitro* ligation-PCR’ approach.

These strategies have several advantages over the commonly used plasmid-based or CRISPR/Cas9-based approaches. First, our strategies for library creation are easy to perform, only requiring PCR reaction, restriction digestion, ligation, and cell transfection, all of which are routine experiments in wet biological laboratories and are fairly easy to master for novices. Second, the screening capacity of libraries constructed is considerable. As an example, the 3NNK-EGFP *in-vitro* ligation-PCR library constructed in two 10-cm dish-seeded HEK293T cells theoretically contains more than 8×10^5^ variant members (20×20×20×100), since the 1:100 diluted library covers all eight X65-Y66-G67-pattern green fluorescence-chromophores **(Figure 4)**, This library size is relatively large from the perspective of screening capacity ^[7, 30, 31]^, especially considering its further growth potential with the expansion of transfection scale. Third, piggyBac transposon system offers a large cargo-carrying capacity (over 200 kb) ^[22]^, comparing to <200 bp insertion mediated by ssODNs in existing CRISPR/Cas9-based strategies. This large cargo-carrying capacity provides the opportunity of simultaneous insertion of drug selection genes into the genome, greatly facilitating the downstream selection of cells with successful genomic integration. And to our best knowledge, we for the first time found the piggyBac transposon system to be with the capability to mediate the genomic integration of *in-vitro* ligation dsDNAs or PCR products with bilateral ITRs. Forth, the *in-vitro* ligation library construction strategy we provide is flexible, and easily adaptive to other transposon systems, e.g. Sleeping Beauty system ^[32]^ and Tol2 system ^[33]^, only requiring the addition of the corresponding transposase recognition sequences on both termini. Recently, large serine recombinases (LSRs) have been found to possess ability to integrate linear PCR amplicons into genomic DNA ^[34, 35]^. Additionally, by including homologous arm sequences, the *in-vitro* ligation dsDNAs or *in-vitro* ligation-PCR products could also serve as the homologous repair template for CRISPR/Cas9-mediated HDR. Thus, the combined application of our *in-vitro* ligation library construction strategy with LSRs or CRISPR-Cas9 system for the establishment of genomic-integrated DMS libraries in mammalian cells is also theoretically feasible.

However, at the same time, we also need to pay attention to some limitations of these novel strategies. First, the piggyBac transposase mediates the transposon integration into ‘TTAA’ sites which disperse in the genome randomly. This may potentially lead to insertion mutagenesis and produce inconsistent expression, and it is also hard to achieve single-copy genomic integration. These shortcomings of piggyBac transposon system seem to have no effect on the positive selection of green fluorescence-generating proteins or transcriptional activator peptides we presented here. In the future study, we plan to borrow CRISPR-Cas9 system to deliver *in-vitro* ligation dsDNAs into the safe harbor of human genome to circumvent these limitations. Second, these strategies are more suitable for the induction of exogenous libraries into mammalian cell genome than for the creation of in situ mutagenesis libraries of endogenous genes of interest. *In-vitro* ligation dsDNAs with homologous arms at the termini in combination with CRISPR-Cas9 system will help breaking through this limitation. Third, the NNK degenerate codons we utilized here induce certain redundancy into the constructed libraries which requires additional screening efforts, although this is already the case after optimization (comparing to NNN codons). Further reduction of the redundancy can rely on the other randomization schemes, e.g. the SILM strategy or 22c-trick, which use the mixtures of ordinary primers to achieve zero or near-zero redundancy ^[30, 36-38]^. However, the economic cost of library construction will increase due to the larger number of primers, which should be traded off by investigators based on their own situations ^[38]^. Forth, in the current strategy design, a non-palindromic restriction enzyme BssSI (also known as BauI) was used to digest the dsDNAs, based on the consideration that the non-palindromic sticky ends block the upstream-to-upstream or downstream-to-downstream inaccurate dsDNA ligation and lead to a higher yield of accurate upstream-to-downstream ligation products. However, on the other hand, commercially available non-palindromic endonucleases are relatively few, and there will be no suitable non-palindromic restriction sites in some sequence context. Gibson Assembly system may help circumventing the restriction enzyme digestion, though further practical attempts are needed to make it a reality.

In the tentative screening of green fluorescence-chromophore sequences performed in this study, we identified eight 3-amino-acid combinations confirming to X65-Y66-G67 pattern (AYG, CYG, GYG, LYG, MYG, SYG, TYG, and VYG) and further verified their green fluorescence-generating ability under 488-nm laser excitation by flow cytometry. Among them, T65-Y66-G67 is the original sequence of EGFP-chromophore, and S65-Y66-G67 is that of wtGFP-chromophore. The fluorescence of wtGFP is believed to depend on the autocatalytic formation of *p*-hydroxybenzylideneimidazolinone chromophore by Ser65, Tyr66 and Gly67, which is buried in the center of a cylindrical geometry formed by an eleven-stranded β-barrel structure playing a protective role ^[25, 26, 28, 39-41]^. The cyclic chromophore formation is proposed to start with the nucleophilic attack by the amide nitrogen of Gly67 on the carbonyl carbon atom of Ser65, and the following dehydration leads to the formation of a cyclized ring. And the subsequent oxidation of Tyr66 is indispensable for the mature GFP chromophore formation and green fluorescence generation ^[25, 28]^. Tsien et al. revealed that wtGFP mutants in which Ser65 is replaced by Thr, Cys, Leu, Val or Ala have spectra with the absorption peaks ranging from 470-490 nm and the emission peak ranging from 504-511 nm, and their amplitudes are greater than that of wtGFP, while the replacement of Ser65 with Arg, Asn, Asp, Phe, or Trp results in dramatic reduction of fluorescence intensity ^[28, 42, 43]^. Our present study perfectly reproduced these phenomena, and further demonstrated another two mutants to be green fluorescence-generating (S65G and S65M). We also found that EGFP-mutants with the replacements of Ser65 by Asp, Glu, Phe, His, Ile, Lys, Asn, Pro, Gln, Arg, Trp or Tyr could not generate green fluorescence under 488-nm laser excitation. Considering that these 12 residues are with relatively big side chain structures, we speculate that this phenomenon may owe to the steric hindrance effects of side chains which may hinder the formation of cyclic chromophore or the peripheric tightly interwoven β-barrel architecture. Nevertheless, it is also worth noting that our present study only detected EGFP-mutants under 488-nm laser excitation by flow cytometry, which may neglect those green fluorescence-generating mutants with shorter or longer excitation wavelength, although this limitation does not affect the interpretability of the resulting measurements of mutational impacts.

The identification of novel transcriptional activator peptide TAP1 well demonstrated the practical feasibility of the ‘piggyBac-*in-vitro* ligation-PCR’ strategy in functional screening in the context of mammalian cells. TAP1 peptide has a characteristic sequence motif DNFDW, in which one hydrophobic residue Phe is flanked by three carbonyl-containing amino acids. The famous transcriptional activator VP16 domain has a closely analogous sequence motif (DAL<u>DDFDL</u>DML). There have been several attempts on resolving the critical structural elements of VP16 domain, which can be used for our reference to study TAP1^[29, 44, 45]^. The negative charge of VP16 domain is considered to be critical to transcriptional activation, and the replacement of increasing number of Asp residues with uncharged residues leads to a progressive decrease of transcriptional activity ^[29]^. TAP1 peptide contains two acidic residues, much less than five in the VP16 domain. But unexpectedly, we observed an increase of transcriptional activity of TAP1 compared to VP16 in a transient transfection system, implying that the absolute value of negative charge may not be the only determinant of the transcriptional activation ability. Schaffiner et a al. has narrowed down the minimal transcriptional activation domain of VP16 to the sequence motif DDFDL ^[44]^. The analogous sequence motif DNFDW in TAP1 retains the Phe residue whose side group is considered to participate in transcriptional activation, while substituting the second Asp in DDFDL with another carbonyl-containing residue Asn and the Leu with another hydrophobic residue Trp **(Figure 6C)**. Considering the prominent position of DDFDL motif in transcriptional activation by VP16, we infer that the sequence motif DNFDW also plays a central role in transcriptional activity of TAP1 peptide. There are multiple hydrophobic residues (<u>LIF</u>TY<u>M</u>DN<u>F</u>D<u>WW</u>, 6 out of 12) in TAP1 peptide. The sequence pattern consisting predominantly of hydrophobic and carbonyl-containing amino acids is a common feature among various transcriptional activators, especially the so-called acidic activators ^[29, 44, 46, 47]^, indicating that the transcriptional activator peptide screened out by our novel approach is in line with the law of natural selection.

In summary, here we developed two novel methodologies to construct genome-integrated DMS libraries in mammalian cells. The enrichment model for green fluorescence-chromophore amino acid sequences demonstrated the feasibility of both strategies in DMS library construction, and the identification of novel transcriptional activator peptides further verified the applicability in functional screening. Our strategies are easy to perform and easily adaptive, help constructing libraries with considerable screening capacity, which may have great application potential in future mammalian cell DMS library construction.

## Acknowledgements

This work was supported by the National Natural Science Foundation of China [31971388 to W.S., 31870860 to S.L.], Tianjin Key Medical Discipline (Specialty) Construction Project [TJYXZDXK-012A to S.L.], and Fundamental Research Funds for the Central Universities [63211138 and 63221425 to W.S.].

SL conceived and supervised the study. SL and WS designed the experiments, analyzed the data and prepared the figures. SL, WS and YF wrote the paper. YW, YZ, YL and KZ performed the experiments and collected the data. All authors discussed the results and reviewed the manuscript. The authors declare that they have no competing interests.

