## Supplementary figure 1 for "piggyBac-Mediated Genomic Integration of Linear dsDNA-Based Library for Deep Mutational Scanning in Mammalian Cells"

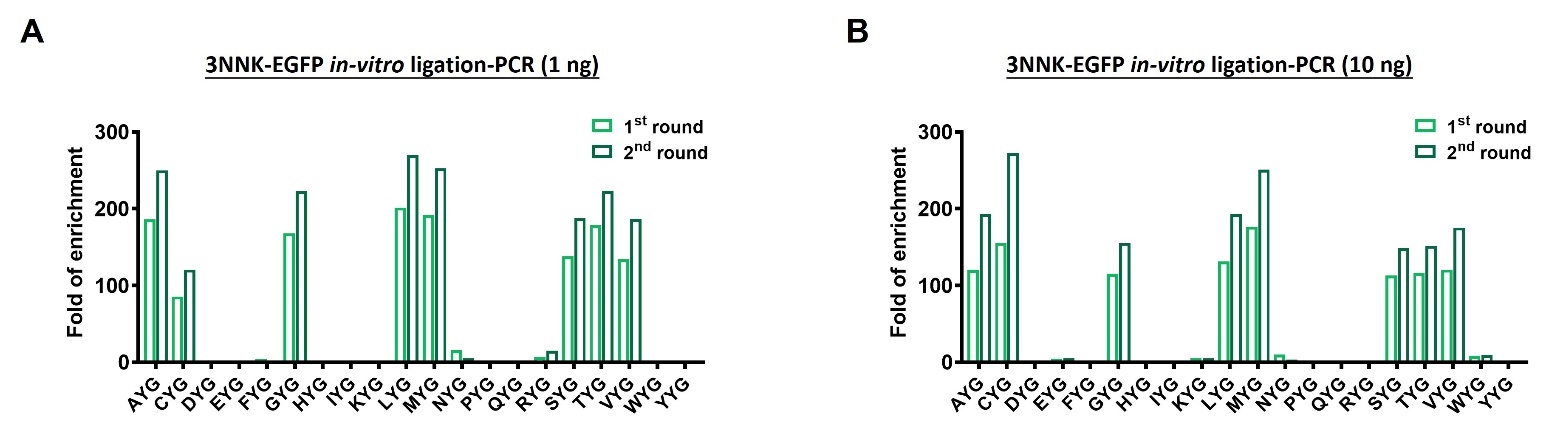

**Supplementary Figure 1. The fold of enrichment of each X65-Y66-G67-pattern 3-amino-acid combination in 3NNK-EGFP *in-vitro* ligation-PCR libraries after each round of FACS sorting.**

**(A)** 3NNK-EGFP *in-vitro* ligation-PCR (1 ng) library. (B) 3NNK-EGFP *in-vitro* ligation-PCR (10 ng) library.
