## Supplementary figure 2 for "piggyBac-Mediated Genomic Integration of Linear dsDNA-Based Library for Deep Mutational Scanning in Mammalian Cells"

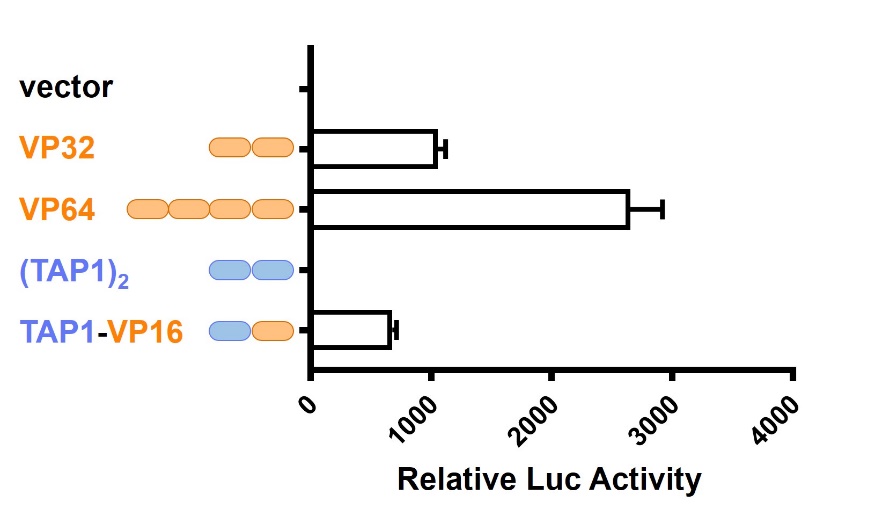

**Supplementary Figure 2. Transcriptional activity of TAP1 derivatives.**

Transcriptional activation potential of (TAP1)_2_ and TAP1-VP16. The coding sequences of (TAP1)_2_ and TAP1-VP16 were cloned into pBIND vector coding GAL4 DNA-binding domain (namely pBIND-XTEN-(TAP1)_2_ and pBIND-XTEN-TAP1-VP16), and were respectively co-transfected with pG5luc vector containing five GAL4 binding sites upstream of a minimal TATA box and luciferase CDS into HEK293T cells. The transcriptional activity of each peptide was assessed by the relative Luc activity (Fluc/Rluc) measured by the dual luciferase reporter assay. All data are displayed as the mean ± SD (n = 4).
