## Supplementary table 1 for "piggyBac-Mediated Genomic Integration of Linear dsDNA-Based Library for Deep Mutational Scanning in Mammalian Cells"

**Supplementary Table 1. PCR templates and primers used in the study.**

| **Amplicon** | **Template** | **Forward primer** | **Reverse primer** | **Related Figures** |
| --- | --- | --- | --- | --- |
| PuroR upstream-dsDNAs | PuroR-template-plasmid | GATTCTGTGGATAACCGTATTACCG | gtcgagcccgacTCTCGTgaggaagagttcttgcagctcggtgac | Figure 1 |
| PuroR downstream-dsDNAs | PuroR-template-plasmid | gaactcttcctcACGAGAgtcgggctcgacatcggcaaggtgtgg | TCTATCAGGGCGATGGCCCACTACG | Figure 1 |
| PuroR PCR products | PuroR-template-plasmid | GATTCTGTGGATAACCGTATTACCG | TCTATCAGGGCGATGGCCCACTACG | Figure 1 |
| EGFP-chromophore-randomized upstream-dsDNAs | GFP-chromophore^del^-library-template-plasmid | GATTCTGTGGATAACCGTATTACCG | TTGTGCCCCAGGATGTTGCCGTCCT | Figure 2 |
| EGFP-chromophore-randomized (2NNK) downstream-dsDNAs | GFP-chromophore^del^-library-template-plasmid | TGGCCCACCCTCGTGACCACCCTGNNKNNKGGCGTGCAGTGCTTCAGCCGCTACCCCGAC | TCTATCAGGGCGATGGCCCACTACG | Figure 2 |
| EGFP-chromophore-randomized (3NNK) downstream-dsDNAs | GFP-chromophore^del^-library-template-plasmid | TGGCCCACCCTCGTGACCACCCTGNNKNNKNNKGTGCAGTGCTTCAGCCGCTACCCCGAC | TCTATCAGGGCGATGGCCCACTACG | Figure 2 |
| EGFP-chromophore-randomized (3NNK) dsDNAs  (PCR-amplified) | EGFP-chromophore-randomized (3NNK) dsDNAs  (by *in-vitro* ligation) | GATTCTGTGGATAACCGTATTACCG | TCTATCAGGGCGATGGCCCACTACG | Figure 3-4 |
| EGFP-chromophore-deleted dsDNAs | GFP-chromophore^del^-library-template-plasmid | GATTCTGTGGATAACCGTATTACCG | TCTATCAGGGCGATGGCCCACTACG | Figure 4 |
| Amplicon for PE-sequencing library construction (for chromophore sequences) | HEK293T library cell genomic DNA | CTACGGCAAGCTGACCCTGAAGTTC | CGCGGGTCTTGTAGTTGCCGTCGTC | Figure 2-4 |
| Transcriptional activator peptide library upstream-dsDNAs | GAL4-XTEN-library-template-plasmid | GATTCTGTGGATAACCGTATTACCG | GACACCTACTCAGACAATGCGATGC | Figure 6 |
| Transcriptional activator peptide library downstream-dsDNAs | GAL4-XTEN-library-template-plasmid | CGCACTCACGAGTGAGTCCGCCACACCCGAATCCNNKNNKNNKNNKNNKNNKNNKNNKNNKNNKNNKNNKTAAACCCAGCTTTCTTGTACAAAGTG | TCTATCAGGGCGATGGCCCACTACG | Figure 6 |
| Transcriptional activator peptide library dsDNAs (PCR-amplified) | Transcriptional activator peptide library dsDNAs  (by *in-vitro* ligation) | GATTCTGTGGATAACCGTATTACCG | TCTATCAGGGCGATGGCCCACTACG | Figure 6 |
| Amplicon for PE-sequencing library construction (for transcriptional activator peptide sequences) | HEK293T library cell genomic DNA | GTTGACTGTATCGCCGGAATTCCCG | GTGGCACCTTCCAGGGTCAAGGAAG | Figure 6 |

Notes: N = T or C or A or G; K = T or G.
