## Supplementary table 2 for "piggyBac-Mediated Genomic Integration of Linear dsDNA-Based Library for Deep Mutational Scanning in Mammalian Cells"

**Supplementary Table 2. Top 10 enriched 2-amino-acid combinations in 2NNK-EGFP *in-vitro* ligation library.**

| **NO.** | **2-amino-acid combinations** | **Initial RPM** | **RPM after 1^st^ selection** | **Enrichment for 1^st^ selection** | **RPM after 2^nd^ selection** | **Enrichment for 2^nd^ selection** |
| --- | --- | --- | --- | --- | --- | --- |
| 1 | MY | 1139.17 | 12125.11 | 10.64 | 15675.13 | 13.76 |
| 2 | AY | 2634.41 | 25423.77 | 9.65 | 30479.93 | 11.57 |
| 3 | TY | 2219.29 | 19162.76 | 8.63 | 24262.65 | 10.93 |
| 4 | GY | 1220.63 | 9169.16 | 7.51 | 12598.56 | 10.32 |
| 5 | SY | 4895.59 | 37417.45 | 7.64 | 50282.49 | 10.27 |
| 6 | VY | 3076.29 | 26137.04 | 8.50 | 31266.93 | 10.16 |
| 7 | LY | 5569.03 | 36319.74 | 6.52 | 44667.70 | 8.02 |
| 8 | AW | 2810.70 | 14388.05 | 5.12 | 18951.42 | 6.74 |
| 9 | CY | 1434.44 | 8183.67 | 5.71 | 8185.43 | 5.71 |
| 10 | IK | 783.69 | 3184.53 | 4.06 | 3987.23 | 5.09 |
