## Supplementary table 3 for "piggyBac-Mediated Genomic Integration of Linear dsDNA-Based Library for Deep Mutational Scanning in Mammalian Cells"

**Supplementary Table 3. Top 15 enriched 3-amino-acid combinations in 3NNK-EGFP *in-vitro* ligation library.**

| **NO.** | **3-amino-acid combinations** | **Initial RPM** | **RPM after 1^st^ selection** | **Enrichment for 1^st^ selection** | **RPM after 2^nd^ selection** | **Enrichment for 2^nd^ selection** |
| --- | --- | --- | --- | --- | --- | --- |
| 1 | GYG | 134.99 | 13901.53 | 102.98 | 66593.33 | 493.32 |
| 2 | DTK | 0.66 | 240.63 | 365.43 | 288.28 | 437.80 |
| 3 | CYG | 113.59 | 7466.41 | 65.73 | 48839.96 | 429.97 |
| 4 | TYG | 158.04 | 12669.19 | 80.17 | 67465.92 | 426.90 |
| 5 | HMT | 52.68 | 3957.43 | 75.12 | 21709.40 | 412.11 |
| 6 | HKM | 5.27 | 707.84 | 134.37 | 1925.67 | 365.55 |
| 7 | AID | 11.85 | 1211.50 | 102.21 | 3432.06 | 289.56 |
| 8 | NTM | 91.20 | 3849.61 | 42.21 | 24088.38 | 264.13 |
| 9 | AYG | 67.49 | 8215.91 | 121.73 | 15606.98 | 231.23 |
| 10 | FRM | 128.40 | 3704.30 | 28.85 | 26396.70 | 205.57 |
| 11 | LYG | 137.62 | 9670.65 | 70.27 | 25582.39 | 185.89 |
| 12 | ISQ | 61.90 | 3066.77 | 49.55 | 11336.89 | 183.16 |
| 13 | FPG | 84.29 | 2996.46 | 35.55 | 14447.15 | 171.41 |
| 14 | AWG | 94.16 | 7697.66 | 81.75 | 15340.36 | 162.91 |
| 15 | TQN | 109.97 | 3461.06 | 31.47 | 17807.52 | 161.93 |
