## Supplementary table 4 for "piggyBac-Mediated Genomic Integration of Linear dsDNA-Based Library for Deep Mutational Scanning in Mammalian Cells"

**Supplementary Table 4. Top 10 enriched 2-amino-acid combinations in 3NNK-BFP *in-vitro* ligation library.**

| **NO.** | **2-amino-acid combinations** | **Initial RPM** | **RPM after 2^nd^ selection** | **Enrichment for 2^nd^ selection** |
| --- | --- | --- | --- | --- |
| 1 | SH | 5800.01 | 215181.25 | 37.10 |
| 2 | NP | 1306.15 | 25291.06 | 19.36 |
| 3 | IT | 2066.86 | 21863.43 | 10.58 |
| 4 | HN | 904.12 | 8287.00 | 9.17 |
| 5 | SD | 3172.29 | 26889.23 | 8.48 |
| 6 | FP | 3043.13 | 25333.89 | 8.32 |
| 7 | HE | 639.40 | 4950.40 | 7.74 |
| 8 | LM | 3588.28 | 27732.93 | 7.73 |
| 9 | DD | 995.76 | 7138.13 | 7.17 |
| 10 | CQ | 1384.69 | 9230.21 | 6.67 |
