## Supplementary table 5 for "piggyBac-Mediated Genomic Integration of Linear dsDNA-Based Library for Deep Mutational Scanning in Mammalian Cells"

**Supplementary Table 5. Top 10 enriched 3-amino-acid combinations in 3NNK-EGFP *in-vitro* ligation-PCR (1 ng) library.**

| **NO.** | **3-amino-acid combinations** | **Initial RPM** | **RPM after 1^st^ selection** | **Enrichment for 1^st^ selection** | **RPM after 2^nd^ selection** | **Enrichment for 2^nd^ selection** |
| --- | --- | --- | --- | --- | --- | --- |
| 1 | LYG | 124.54 | 25063.23 | 201.24 | 33611.51 | 269.88 |
| 2 | MYG | 42.55 | 8161.66 | 191.81 | 10741.69 | 252.45 |
| 3 | AYG | 82.47 | 15342.39 | 186.03 | 20625.46 | 250.09 |
| 4 | TYG | 68.13 | 12163.49 | 178.54 | 15209.13 | 223.24 |
| 5 | GYG | 81.04 | 13618.17 | 168.05 | 18028.65 | 222.47 |
| 6 | SYG | 124.78 | 17223.61 | 138.03 | 23395.82 | 187.49 |
| 7 | VYG | 129.32 | 17319.57 | 133.92 | 24134.91 | 186.62 |
| 8 | NQF | 0.24 | 26.88 | 112.44 | 31.92 | 133.55 |
| 9 | CYG | 76.26 | 6532.64 | 85.67 | 9189.03 | 120.50 |
| 10 | QHK | 2.63 | 171.07 | 65.06 | 227.02 | 86.34 |
