## Supplementary table 6 for "piggyBac-Mediated Genomic Integration of Linear dsDNA-Based Library for Deep Mutational Scanning in Mammalian Cells"

**Supplementary Table 6. Top 10 enriched 3-amino-acid combinations in 3NNK-EGFP *in-vitro* ligation-PCR (10 ng) library.**

| **NO.** | **3-amino-acid combinations** | **Initial RPM** | **RPM after 1^st^ selection** | **Enrichment for 1^st^ selection** | **RPM after 2^nd^ selection** | **Enrichment for 2^nd^ selection** |
| --- | --- | --- | --- | --- | --- | --- |
| 1 | CYG | 54.78 | 8516.90 | 155.48 | 14931.82 | 272.58 |
| 2 | MYG | 24.94 | 4401.30 | 176.46 | 6253.18 | 250.71 |
| 3 | AYG | 121.45 | 14580.03 | 120.05 | 23443.15 | 193.03 |
| 4 | LYG | 163.17 | 21425.87 | 131.31 | 31426.11 | 192.59 |
| 5 | VYG | 140.10 | 16890.67 | 120.56 | 24561.86 | 175.32 |
| 6 | YYN | 0.23 | 13.85 | 59.42 | 37.12 | 159.23 |
| 7 | GYG | 109.33 | 12586.33 | 115.13 | 16996.74 | 155.47 |
| 8 | TYG | 65.97 | 7670.64 | 116.28 | 9997.67 | 151.55 |
| 9 | SYG | 171.10 | 19296.10 | 112.78 | 25390.99 | 148.40 |
| 10 | EFI | 8.62 | 450.83 | 52.27 | 755.17 | 87.56 |
